# Crustacean cardiac ganglion model reveals constraints on morphology and conductances

**DOI:** 10.1101/2021.10.21.464612

**Authors:** Dan Dopp, Pranit Samarth, Jing Wang, Daniel Kick, David J. Schulz, Satish S. Nair

## Abstract

The crustacean cardiac ganglion (CG) network coordinates the rhythmic contractions of the heart muscle to control the circulation of blood. The network consists of 9 cells, 5 large motor cells (LC1-5) and 4 small endogenous pacemaker cells (SCs). We report a new three-compartmental biophysical model of an LC that is morphologically realistic and includes provision for inputs from the SCs via a gap-junction coupled spike-initiation-zone (SIZ) compartments. To determine physiologically viable LC models in this realistic configuration, maximal conductances in three compartments of an LC are determined by random sampling from a biologically-characterized 9D-parameter space, followed by a three-stage rejection protocol that checks for conformity with electrophysiological features from single cell traces. LC models that pass the single cell rejection protocol are then incorporated into a network model which is then used in a final rejection protocol stage. Using disparate experimental data, the study provides hitherto unknown structure-function insights related to the crustacean cardiac ganglion large cell, including predictions about morphology including the role of its SIZ, and the differential roles of active conductances in the three compartments. Further, we extend analyses of emergent conductance relationships and correlations in model neurons relative to their biological counterparts, allowing us to make inferences both with respect to the biological system as well as the implications of the ability to detect such relationships in populations of model neurons going forward.

## INTRODUCTION

Neurons are endowed with a rich and complex set of intrinsic and synaptic conductances that control their electrical activity [1; 2]. Although the role of such a varied set of conductances is not fully understood, it is natural to expect that neurons of the same cell type would possess similar membrane properties, especially within the same animal. However, experimental findings suggest that maximal conductance levels of individual currents can vary two- to six-fold among same cell types, even within the same animal [3-7] and that different combinations of conductances preserve activity at the single cell level [8; 9]. Computational modeling continues to shed light on the role of such conductance variations in conserving cellular output such as spike and burst characteristics [2; 10-18]. For instance, Prinz et al. [2] explored the maximal conductance space of a single-compartment model neuron to quantify the numerous types of spiking and bursting models and showed that similar patterns of activity could be produced by many different parameter sets, both for single neurons [2] and within small networks [19].

Beyond the broad range of conductance combinations that are associated with convergent outputs among neurons of the same type, there is also substantial reports that among populations of neurons different sets of ionic conductances [5] and ion channel mRNA levels [7; 20] can be correlated with one another in different classes of identified neurons. This suggests that an on-going, rather than developmentally fixed, regulation of specific sets of conductances may be necessary to provide stable output of neurons and networks over the lifetime of an animal. These correlated mRNA and conductance levels can arise from a relatively simple set of feedback control algorithms in computational models [21]. However, there have been few studies that directly demonstrate that these conductance or mRNA relationships are *necessary* to generate appropriate, ongoing neuronal activity in biological neurons. Indeed, compelling computational work has demonstrated that – at least theoretically – such relationships are not necessary to generate robust output in a population of model neurons. For example, previous studies have demonstrated that using a model selection methodology focused on single-cell output in a multi-compartment model results in a population of cells with only weak or no correlations among conductances [14]. However, most studies to date have focused on selecting models based on isolated neuron activity. Therefore, to further extend these analyses, we performed multiple levels of selection that included generating model networks with multiple neurons of a given type, and only selecting cells that perform within biological parameters of the full network output for inclusion in our population. We then looked for conductance correlation relationships among the populations of neurons in our simulated model networks.

For this work, we use a computational model of the crustacean cardiac ganglion (CG) network, based on the crab, *Cancer borealis*. This simple network consists of 4 pacemaker neurons and 5 Large Cell (LC) motor neurons that innervate the heart muscle. The present study extends previous computational investigations that focused on single compartment cardiac ganglion LCs [15; 16; 19] by considering the potential role of conductances in multiple compartments of an LC on its output. Specifically, we develop a new morphologically realistic three-compartment LC model of *Cancer borealis* that incorporates an SIZ compartment and first-hand biological data and validate the model. We then use it to investigate how the distribution as well as potential covariations of conductances affect soma membrane potential response (output). Using a rejection sampling approach with a 9-D parameter space of maximal conductances, we report, as in previous studies, an unbiased approach to determine the role of various conductances in shaping cellular and network function. We extend previous studies by performing model neuron selection in complete CG networks, with constraints drawn from intact network activity as well as single cell electrophysiology and current response data. A population of model LCs generated by such an approach then provided predictions related to the differential roles of conductances in the soma vs. neurite in shaping neuronal output. Further, this population of neurons allowed us to look for emergent conductance relationships both within and across these compartments. Finally, we compare conductance relationships uncovered through the model selection process with a comprehensive set of single-cell mRNA relationships for channels underlying these membrane conductance relationships.

## RESULTS

### Morphologically realistic LC model and SC stimulus reproduces experimental profiles

Building on our previous two-compartment LC model [15; 16; 19], we added morphological realism to the LC by adding a third compartment, the spike initiation zone (SIZ; Fig. 1A). We first matched passive properties (see methods) and waveform data from intact cells LC3 and LC4 (Fig. 1A; [19]). We note that among the five LCs, only LC3-5 are easily accessible, and that LC4 is gap-junction coupled strongly to LC5 [22]. Accordingly, our biological data are from LCs 3-5, while our model predicts network performance for all five LCs, assuming LCs1-2 have similar gap junction coupling as LC4-5 (Fig. 1A).

**Figure 1.**
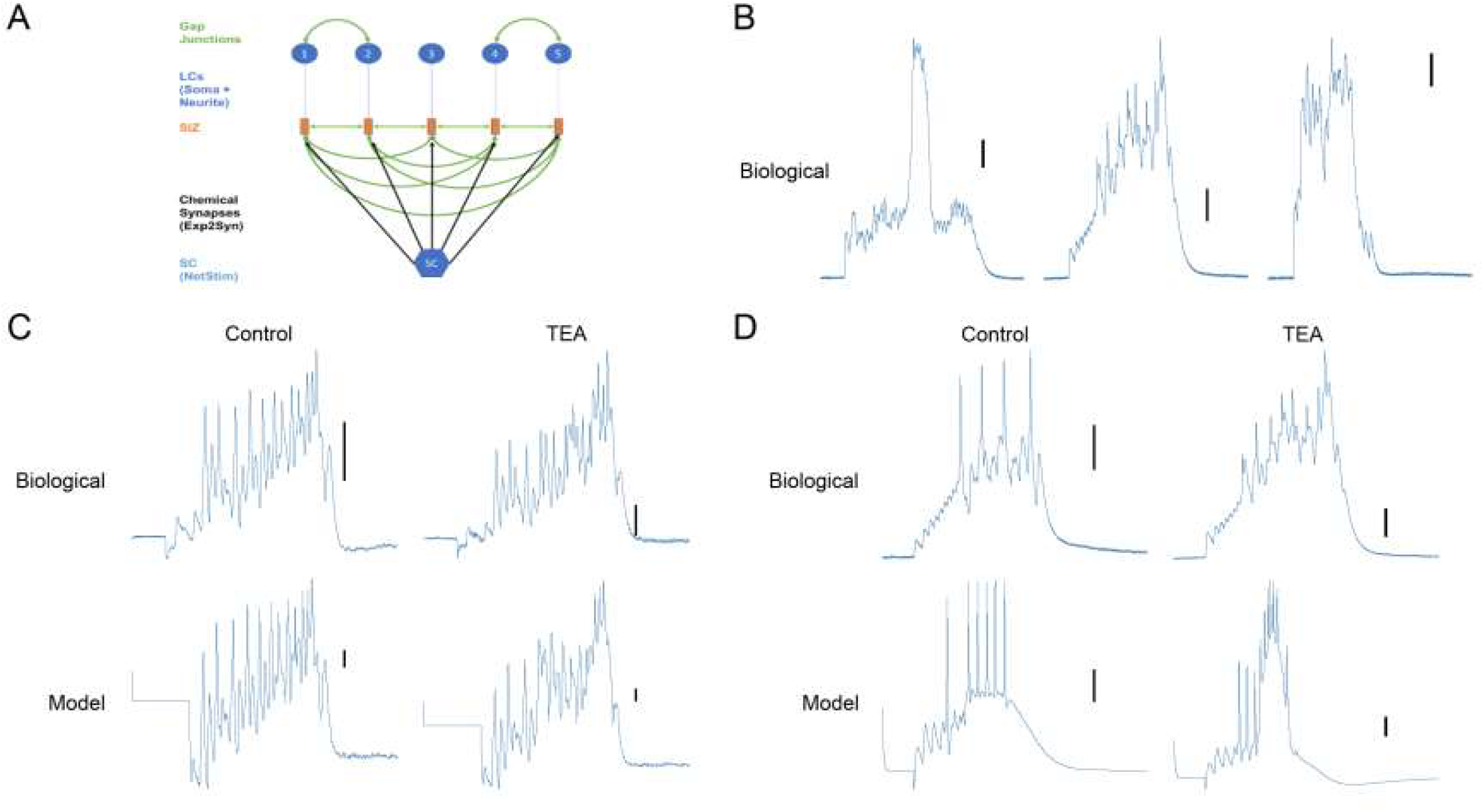
First model of intact LC reproduces experimental data. (A) Model network. Gap Junctions are formed between LC1-2 and LC4-5. Gap Junctions modeled between SIZs using the same fixed conductance value with SC input to each SIZ. (B) Experimental stimulus protocol response of ligated cell in control and TEA conditions -from Jing’s work. Corresponding model responses are shown on the right. (C) Same plots as in panel B, but for intact cells; (D) Variability in experimental TEA responses from our lab.

#### Developing rejection protocol criteria from experimental data

We briefly describe the key characteristics of a modified version of our previous rejection protocol [19] to select model LCs that match experimental data. In the first stage, we sampled a 9-D parameter space of maximal conductances to generate a pool of ligated LCs (soma + neurite with passive conductances) for which the passive properties of resting potential and input resistance were within biological ranges measured in our Lab [22]; this relaxed some of the constraints in our previous protocol (see methods) and permitted more cells to pass this stage. To the cells that passed, we then added active conductances (picked randomly – see below) in the neurite and an SIZ compartment (with fixed conductances) in stage 2, and then provided synaptic input from the SC between the experimentally observed range of 16-32 Hz and retained only cells that had at least one spike with the SC input. This ensured that stage 2 did not pass cells that only had membrane depolarizations but no spikes, reducing the load on the computationally intensive stage 3. In a third stage, we combine five ‘viable’ (passing selection criteria) randomly-selected single cells into a network. The network itself is deemed viable if it satisfied the two experimental observations related to LC3 and LC5: (i) a synchrony value between LC3 and LC5 >0.95 in control (Pearson’s R-squared); and (ii) synchrony between LC3 and LC5 <0.89 in TEA. Results from each of these stages are discussed next.

#### Membrane potential responses of ligated cells

The ligated cell (soma + neurite) model was tested for passive properties based on experimental data from our Lab [22]. From among 150,000 ligated model cells with random conductances selected from the 9-D parameter space (see methods), 100,000 passed stage 1 of the rejection protocol. Figure 1B shows a representative experimental (Fig1B1 & Fig1B2, Pre & Post) and model (Fig1B3 & Fig 1B4, Pre & Post) membrane potential responses of a ligated cell to an experimentally determined input (termed ‘stimulus protocol’; [22]; see methods) in control and post-TEA conditions. After finding viable ligated cells (soma and passive neurite), we attached an SIZ to each cell in the pool for further testing in stages 2 and 3. For this, we designed an SC input to represent experimentally determined SC input characteristics as described next.

#### Designing the SC input to the LC network

In the crustacean, the five-cell LC network receives input from a cluster of four small cells, and we assumed that all five LCs receive a common synchronized input from this SC cell cluster. We designed the SC input as a spike train to a synapse on the SIZ, the properties of which are tuned to mimic experimental voltage responses [23]. Experimental recordings from the Schulz Lab (unpublished data) showed that SC input frequency varied between 16 and 32 Hz in 1 Hz intervals. Also, two components were noted in the experimental data over a typical period of 1000 ms, a steady one that continued over the entire duration, and a second one that lasted for 600 ms, starting from 300 ms and ending at 900 ms. As described in methods, we designed the SC input with the two components, after randomly picking a frequency within the rage of 16-32 Hz.

#### Network Responses - matching responses of intact single cells

The SC input we designed was then used in the next stage to provide input to intact cells formed by adding an SIZ and synapse to the ligated single cell model. Figure 1C1 (control)shows the experimental recording for an intact LC4 cell A typical corresponding response from the model intact cell is shown in Figure 1C3 (PreTEA) and Figure 1C4 (PostTEA),. To make the analysis tractable, we considered the case where the intact cells in a network did not receive input from the other four LCs, i.e., all gap junction coupling between the LCs were disconnected. We consider the SIZ gap-junction coupled case in a later section.

For such an intact single LCs, we initially assumed a passive neurite, i.e., only leak conductance in the neurite. So, to the model cells that passed stage 1, we connected an SIZ and synapse, and used the SC spike train input described in the previous section. Interestingly, none of the 100,000 cells passing stage 1 were able to pass stage 2. This was because the cells had a spiking frequency above 8 Hz in control and did not exhibit a TEA response. However, the SIZ responses did match biological reports. Specifically, the model membrane potential responses at the soma had a depolarization of 10 mV for 1000 ms, and with spikes on top of the depolarization that reached 20 mV in height. This response matched the soma membrane potential response characteristics from our lab that had a depolarization bump of 10 mV for 1000 ms, and spike height of 15 mV (Fig. 1C1). Additionally, the model SIZ spike height attenuated by a factor of 3 (30 to 10 mV) at the soma (figure 5C), matching the corresponding biological recordings from our Lab which had an attenuation factor of 3.66 (55 to 15 mV) (unpublished data; also matched experimental SIZ recording in [23]).

The functional reason for the cells failing in stage 2 was determined to be the excessive leak through the neurite, i.e., although sufficient current entered the neurite from the SIZ, this leakage diminished the amount that reached the soma for raising the response in TEA. Reducing the diameter, and therefore surface area, was found to decrease leakage. However, the neurite diameter had to be 5 μm, which was unrealistically small compared to biological minimum diameter of 10 μm. As a next step, we considered active conductances in the neurite.

### Active conductances necessary in neurite to reproduce experimental network responses

The presence or role of active conductances in the neurite of the CG LC is unknown. For instance, although our mRNA studies suggest the presence of Nap in the LC), the morphological specificity of location is unknown. Similarly, although Ca2+ currents are thought to be present in the neurite, their localization is not fully understood.

#### Role of individual conductances

We adopted a systematic procedure to determine a parsimonious set of active conductances in the neurite. For this, we first inserted only Nap in the neurite, and this helped counter the excessive leakage cited in the previous section, enabling the soma to elicit a TEA response with spiking in SIZ (Fig. 4 A-B Supplemental). With Nap in the neurite, for every 1000 cells that passed stage 1, about 35 passed stage 2. Fewer cells passed because the spike height and LC spike frequency were both found to exceed the upper bounds in the control case. For instance, for a case with SC frequency of 22 Hz (low in experimental traces of SC spikes), the soma spike height was above the upper bound of 30 mV (Table 6) as was the spike frequency, in many of the control cases. The reason for this was that the cell was already close to excitable in the control case with the passive neurite. The key attribute that Nap provided was a TEA response that met the requirements of increased spike frequency and amplitude (Fig S4B) in nearly all cells. So, we considered current channels to reduce the depolarization caused by Nap in the control case while retaining the TEA response. This led to the addition of I_BKKCa but that worked only for some models (Fig S4C), even with maximal conductance of I_BKKCa exceeding the upper bound (Table 2). Furthermore, this manipulation did not provide the variability in TEA responses seen in experimental traces (Fig.1D1-3). Since I_Nap by itself was not sufficient, we then explored whether I_CaT and I_CaS channels could substitute for I_Nap. Even with values of conductances beyond the upper bounds for CaS and CAT channels, the TEA response was inadequate (Fig 4SD), and the peak spike height in the control case was also too high. So, we added I_BKKCa to this set of I_CaT and I_CaS, without Nap. Although this reduced the peak spike height to within permissible ranges, the TEA response was still inadequate (Fig S4E).

**Table 1.**
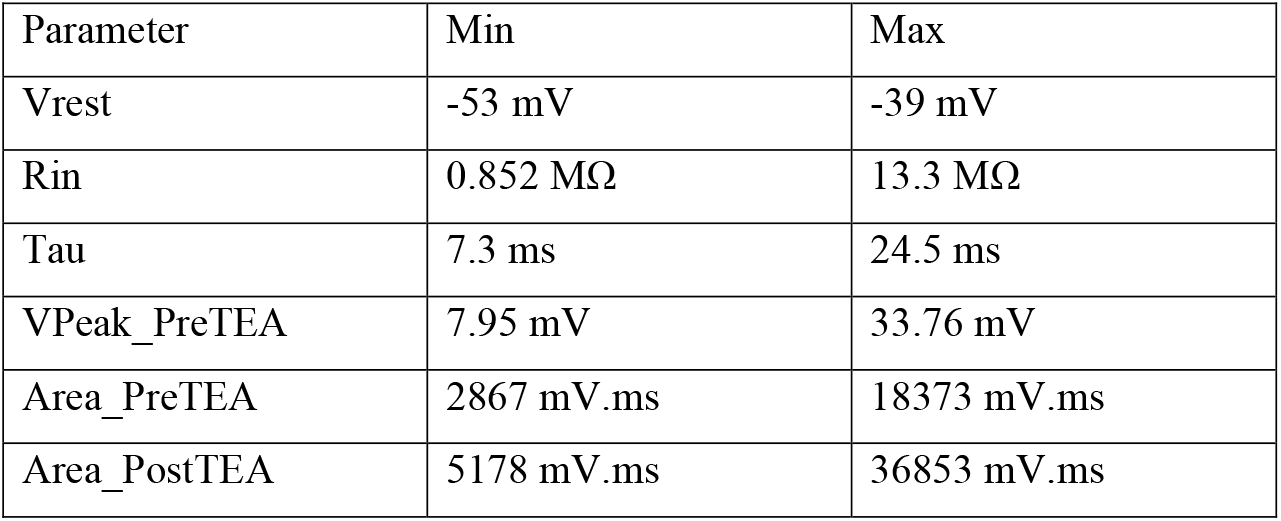
Ranges of properties for selecting valid LCs

**Table 2.** Maximal conductance ranges

| Current | G min (S/cm<sup>2</sup>) | G max (S/cm<sup>2</sup>) |
| --- | --- | --- |
| CaT | 0.00016 | 0.00031 |
| CaS | 6.50E-05 | 0.00013 |
| CAN | 7.00E-05 | 1.50E-04 |
| NaP | 3.50E-05 | 2.30E-04 |
| Leak (All Segments) | 6.20E-05 | 9.70E-04 |
| KA | 0.000172 | 0.0019 |
| Kd (Kd1) | 0.000165 | 0.00127 |
| KCa | 0.00079 | 0.0061 |
| SKKCa | 0.00088 | 0.002 |
| Kd2 | 0.000091 | 0.0005 |

**Table 3.** Ranges of connective and input parameters used in Stage 2&3 rejection

| Parameter | Min | Max |
| --- | --- | --- |
| Exp2Syn Tau1 | 10 ms |  |
| Exp2Syn Tau2 | 60 ms |  |
| Exp2Syn reversal potential | 0 mV |  |
| NetCon weight | 0.01 | 0.1 |
| NetStim spiking frequency | 16Hz | 32Hz |
| NetStime noise level | 0.8 |  |
| LocalGap R | 1 MΩ | |
| InterGap R | 1MΩ | 15 MΩ |

**Table 4.**
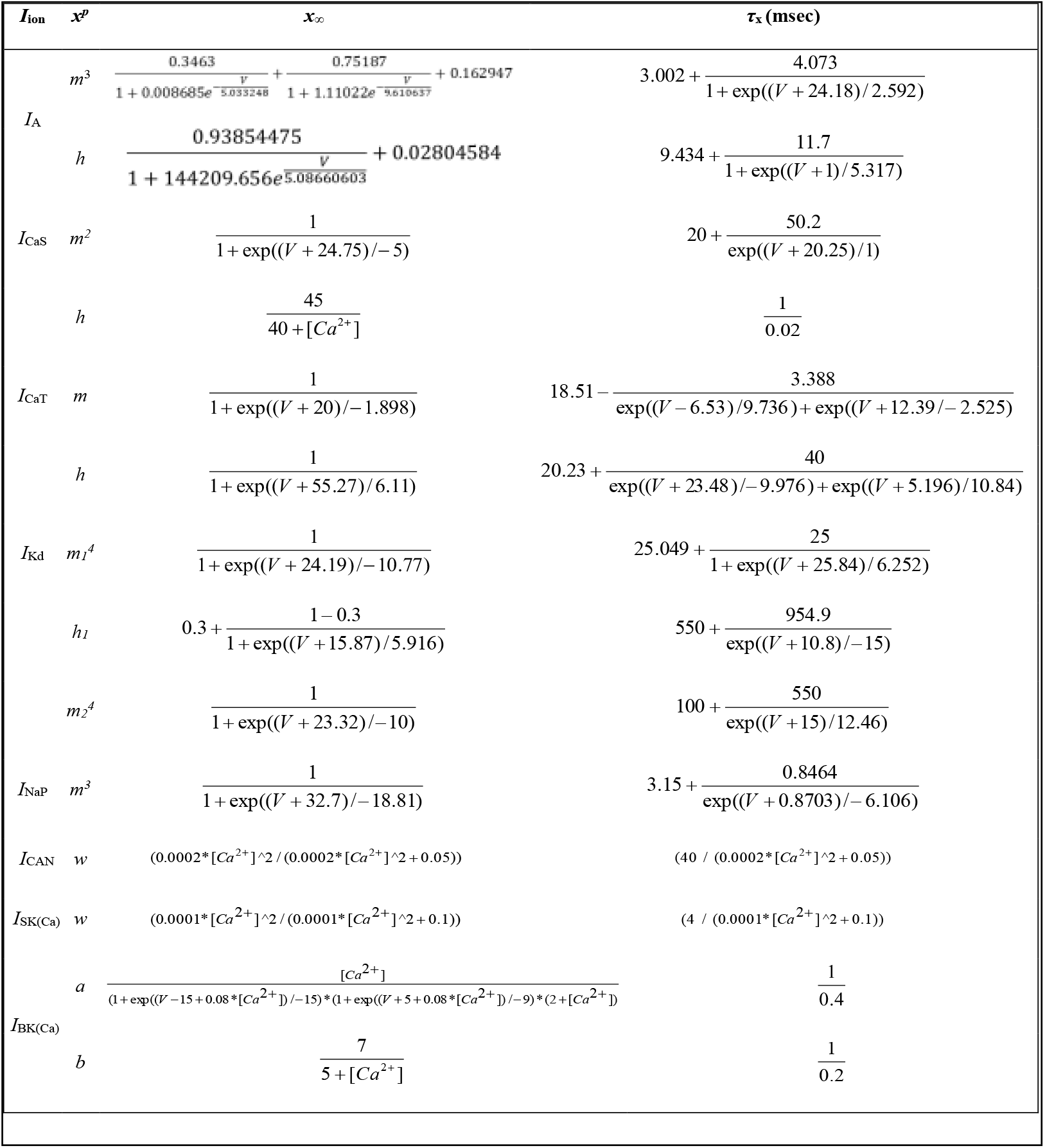
Model Current Parameters

**Table 5.** Ranges of waveform properties for selecting valid intact cells

| Parameter | Min | Max |
| --- | --- | --- |
| Spike number per burst | 4 | 8 |
| Avg LC spiking frequency | 4 Hz | 8 Hz |
| VPeak_PreTEA | 7 mV | 30 mV |
| Area_PreTEA | 2867 mV.ms | 18373 mV.ms |
| Spike number per burst Post TEA | >1.13 x PreTEA<br>SPB | none |
| AVG LC spiking frequency | >1.21x PreTEA<br>frequency | none |
| VPeak Post TEA | >1.3x PreTEA<br>Vpeak | none |
| R<sup>2</sup> synchrony LC3 to LC5 control | 0.95 | 1.0 |
| R<sup>2</sup> synchrony LC3 to LC5 in TEA | 0 | 0.89 |

**Table 6.**
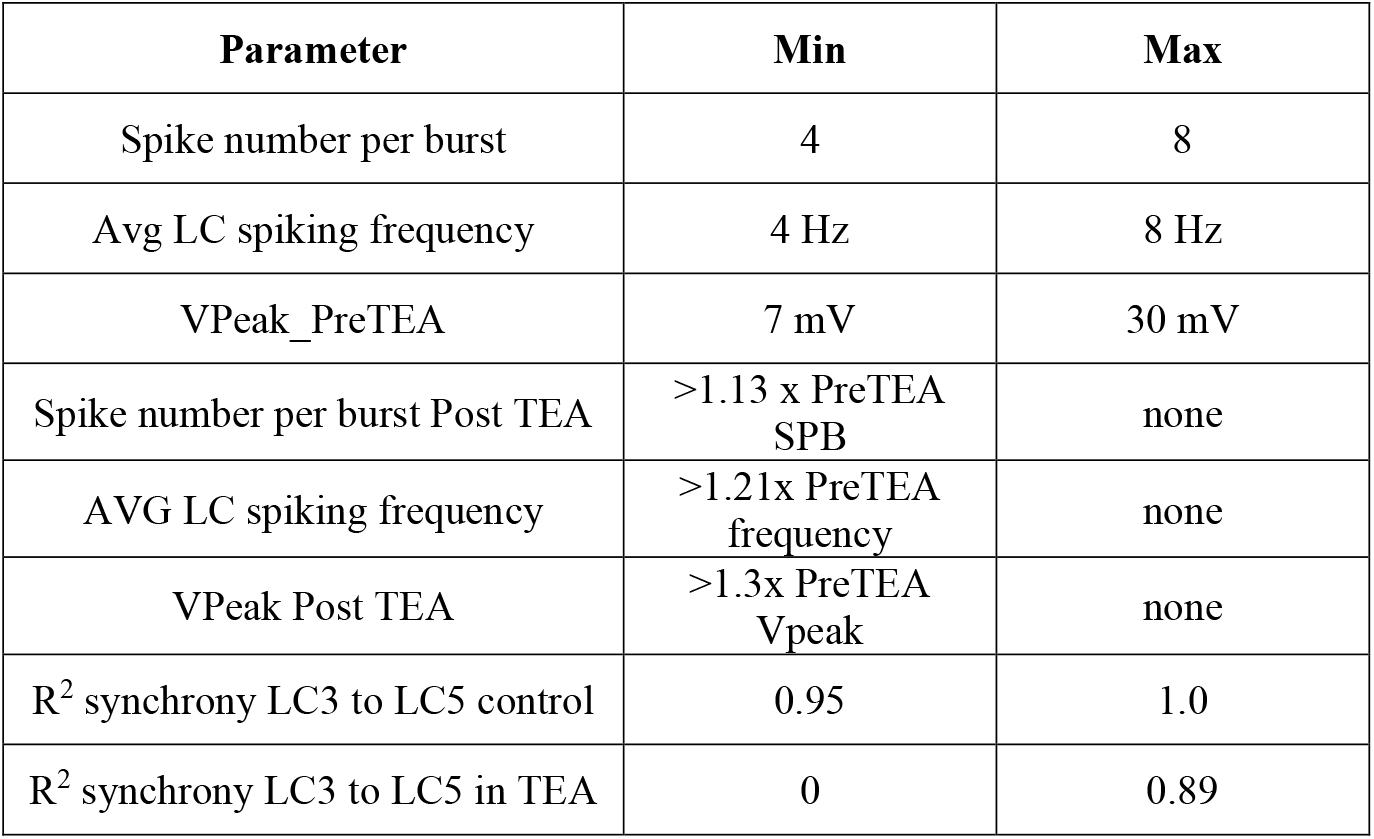
Ranges of waveform and synchrony properties for selecting valid networks

To gain insights into the process, we investigated the mechanism by which I_BKKCa improved the TEA response together with I_Nap (Fig S4C), and produced variable TEA responses seen in experiments, without disturbing the control responses. First, we found that I_BKKCa helped reduce the spike amplitude (compare Fig. S4F and S4G). However, there was little variability in TEA response waveforms of different cells. To explore why, we investigated the underlying current waveforms (Fig 2A). For this cell, I_CaT and I_CaS in the neurite produced less spiking and depolarization in the soma during TEA compared to I_Nap in the neurite (Fig 2 B). The waveform of I_BKKCa corresponds closely in time with those of leak and Nap currents, all of which also closely follow the voltage waveform. This suggests that I_BKKCa has a greater impact on the voltage waveform than did I_CaT and I_CaS. However, the slow wave amplitude could, in general, be modified by I_CaT and I_CaS, allowing the model to exhibit varied TEA waveforms. With all these channels present, the waveform criteria for both control and TEA cases were met by larger numbers of cells (Fig 2B), and there was greater variability in the range of TEA responses. A typical set of TEA responses for five intact cells with all channels present is shown in Fig S4I. In summary, I_BKKCa (with I_CaT and I_CaS) reduced peak spike height and frequency in the control case. Although I_BKKCa did reduce the increased control response caused by I_Nap, it did not affect I_Nap’s facilitation of the TEA response. Furthermore, CaS and CaT channels were found to be important for the generation of varied TEA responses.

**Figure 2.**
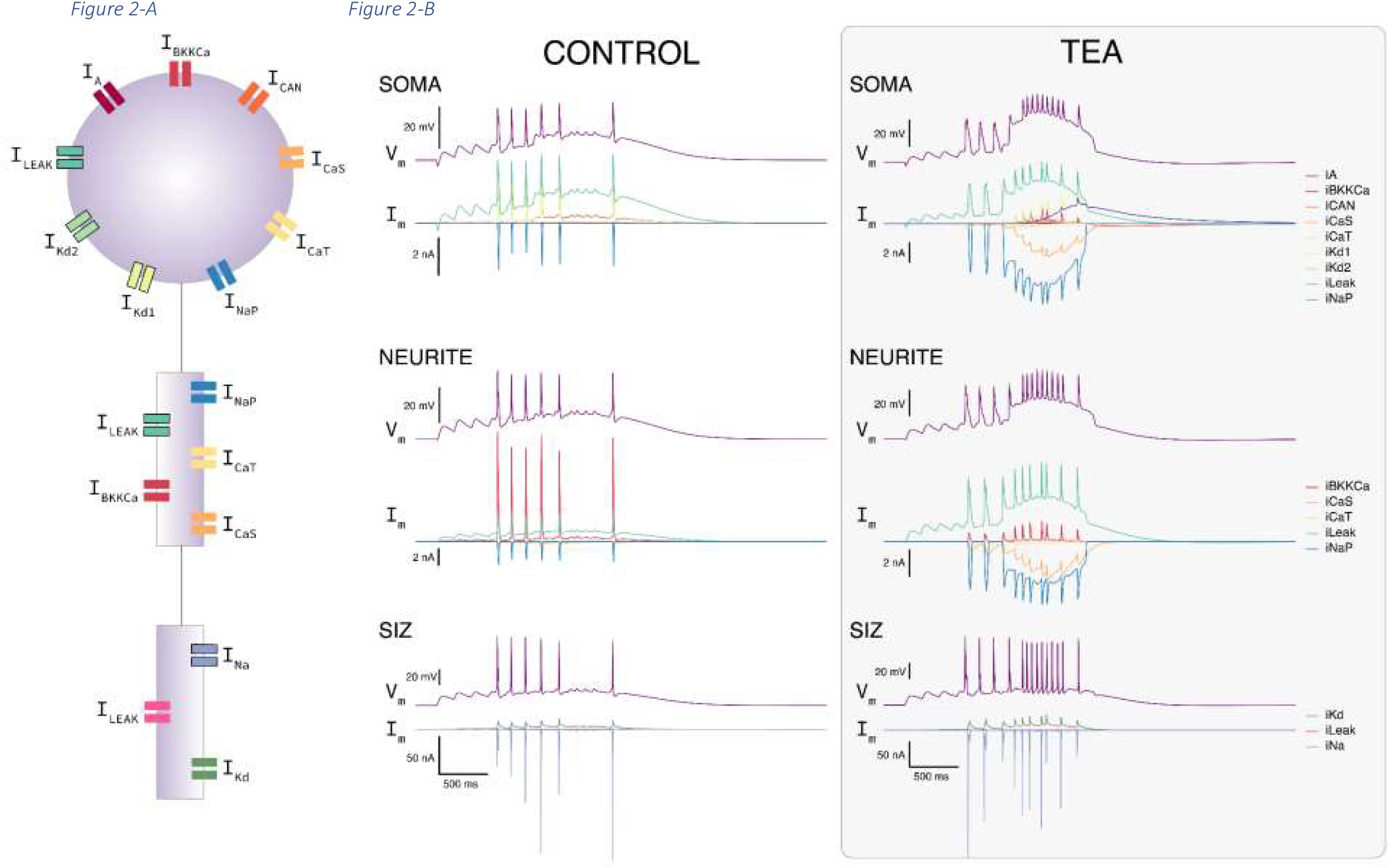
Model predicts presence and roles of active conductances in neurite. (J) Current traces. Top: left, middle right: stimulus protocol as used in the biology lab, Neurite current with soma detached and all currents present, Neurite current with soma detached and only leak current present. Bottom: left, right: Neurite current of intact cell with all currents present, Neurite current of intact cell with only leak present. All model traces are from a cell in a network.

#### Random sampling of neurite conductances and validation checks

Based on the systematic initial trial-and-error investigation of the role of conductances in the neurite discussed in the previous section, we decided to add the following current channels to the neurite using random sampling: CaS, CaT, NaP, BKKCa. The reader is reminded that maximal conductances of the soma currents were finalized in stage 1, and so stage 2 considers only the random selection of maximal conductances for the channels added to the neurite, i.e., stage 2 rejection protocol focused only on conductances in the neurite. Of the 100,000 intact cells that passed stage 1, an active neurite resulted in the total number passing stage 2 to increase to 18,000.

*Validation check of ‘stimulus protocol’ waveform in Ransdell et al. [24]*. In our prior experiments with the ligated soma, Ransdell et al.[24] recorded from intact networks and developed a current trace termed ‘stimulus protocol’ that, when injected into a ligated soma, resulted in a membrane potential profile that matched those from intact network recordings. As a validation experiment, we found that the model current entering the soma from the neurite mimicked the ‘stimulus protocol’ waveform with active, but not a passive dendrite (Fig.S4I).

### Network responses with gap-junction coupling among SIZs

The cells from a sample network that passed this final rejection criteria involving networks (Stage 3) are shown in figure 3 (right panel, top right). Of the 2,000 model cells that passed stage two, a total of 750 passed stage 3 (150 networks). The dissimilar individual responses of LCs to the SC frequency of 18 Hz became highly synchronized when placed in a network with gap-junction connectivity among all LCs at the SIZ and among specific cell pairs LCs1-2 and LCs3-4 at the soma (Fig. 5A1,2). The reader is reminded that a pronounced TEA response is a rejection criterion used in Stages 2 and 3 (see methods). We explored the mechanism by which the gap-junction coupling between soma compartments of LCs1-2 and LCs4-5, and between the SIZ compartments of all cells (Fig. 1A) ensured synchrony among the LCs in the network.

**Figure 3.**
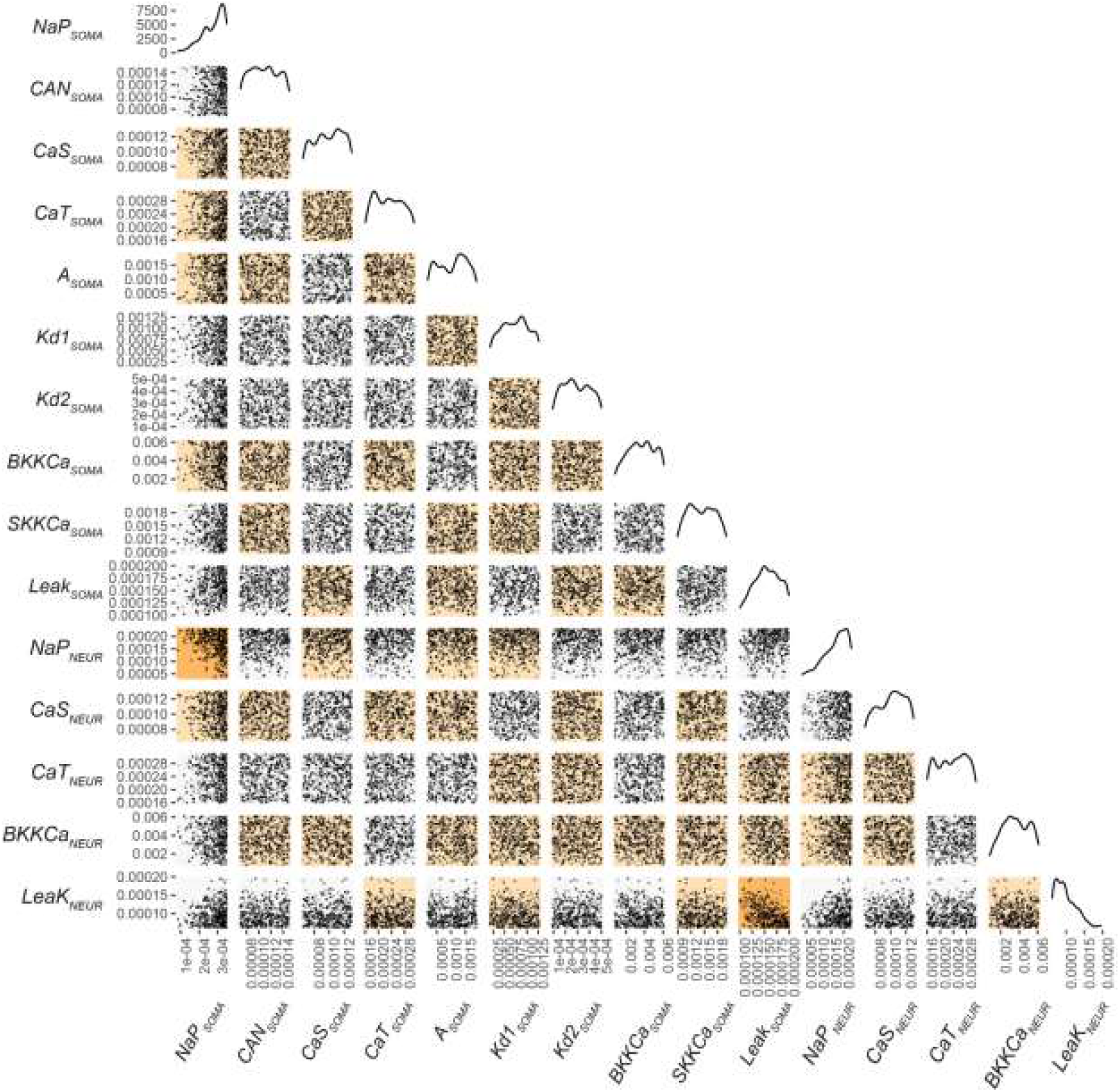
Scatterplots for pairwise relationships among soma, neurite, and across compartment conductances in models of selection level 3. Each dot represents a single model and its values for a given pair of conductances. Along the diagonal are curves that represent the distribution of values for a given conductance as keyed along the bottom axis. Stronger correlations are noted by the increasing intensity of background color for each pairwise relationship.

For this we considered the cells in the example network of Figure 5A2 that passed stage 3 of the rejection protocol, with the same SC input of 18 Hz used for each cell in Fig. 5A. In this network, we found that the gap junction current between soma compartments of LCs1-2 or between LCs4-5 was seven-to ten-fold smaller in magnitude than the synaptic current due to the SC drive. Focusing on one cell, LC1, Fig. 5B1 provides a comparison of the sum of the gap junction current from soma of LC2 to soma of LC1 and from the various SIZ compartments to the SIZ of LC1, labeled as ‘gap junction current’, to the synaptic current into the SIZ of LC1 due to SC input spikes. As can be seen from the traces, the magnitude of the total gap junction current into LC1 was found to be more than seven-fold smaller than that of the synaptic current into LC1. Also, a comparison of traces in Fig. 5A2 and B1 shows that the soma membrane potential response of LC1 is primarily due to the input from SC rather than from input via gap junctions. The total gap junction current is also phase shifted compared to the synaptic current. This sheds light on the role of the synaptic current which is to raise the SIZ membrane potential to spike threshold in ∼500 ms (Fig. 5B1). Once the threshold is reached in the SIZ and spiking is initiated, the gap junction currents ensure that the spikes are synchronized among the cells, and so there is a phase shift of ∼50 ms between the peak of the depolarization due to the synaptic current and the first peak in gap junction current profiles. In summary, the analysis predicts a delineation of the primary functions of the two current types: synaptic (to increase SIZ and soma membrane potential to threshold) and gap junction (to synchronize spikes) and quantifies their magnitudes.

We also note that the observation of the gap junction current being considerably smaller than the synaptic current for each cell justifies the use of intact single cells in stage 2 of the rejection protocol, without considering the gap-junction interaction effects from other cells and provides a *validation check*.

### Conductance parameter space variations following three levels of selection on model neurons

Following each of the stages of selection, we examined the overall distribution of membrane conductances in the soma and neurite compartments in the viable/passing neurons to determine whether each selection criterion limited any conductance to a particular portion of the parameter space (Figure 4). We visualize this in two distinct ways. Figure 4A uses a density plot to describe the conductances that passed a given level of selection. As selection continued, some conductances were more and more limited to a portion of the parameter space, while others maintained a broader range of viable conductances. For example, selection stages 2 and 3 result in a restricted range of distribution of NaP_SOMA_, NaP_NEURITE_, Leak_SOMA_, and Leak_NEURITE_, and Bkkca and Leak. Further, as Vrest was a free parameter in the selection process, we see the strongest selection pressure on this feature, where selection levels 2 and 3 result in a narrow distribution of acceptable values near the high end of the range (Figure 4A). The other active conductances maintained a broad range of acceptable values through all levels of selection. However, these are not normally distributed. Rather, multiple peaks around conductance ranges that enriched for successful models can be seen in most of the conductances (Figure 4A). Finally, because Figure 4A scales each level of selection independently to maximize the opportunity to see variations in the range of conductance values after each selection level, we have also plotted these data in a nested format using a stacked density plot (Figure 4B). This provides the opportunity to see the full parameter space (level 0, purple) and then each round of selection as a subset of the remaining parameter space.

**Figure 4.**
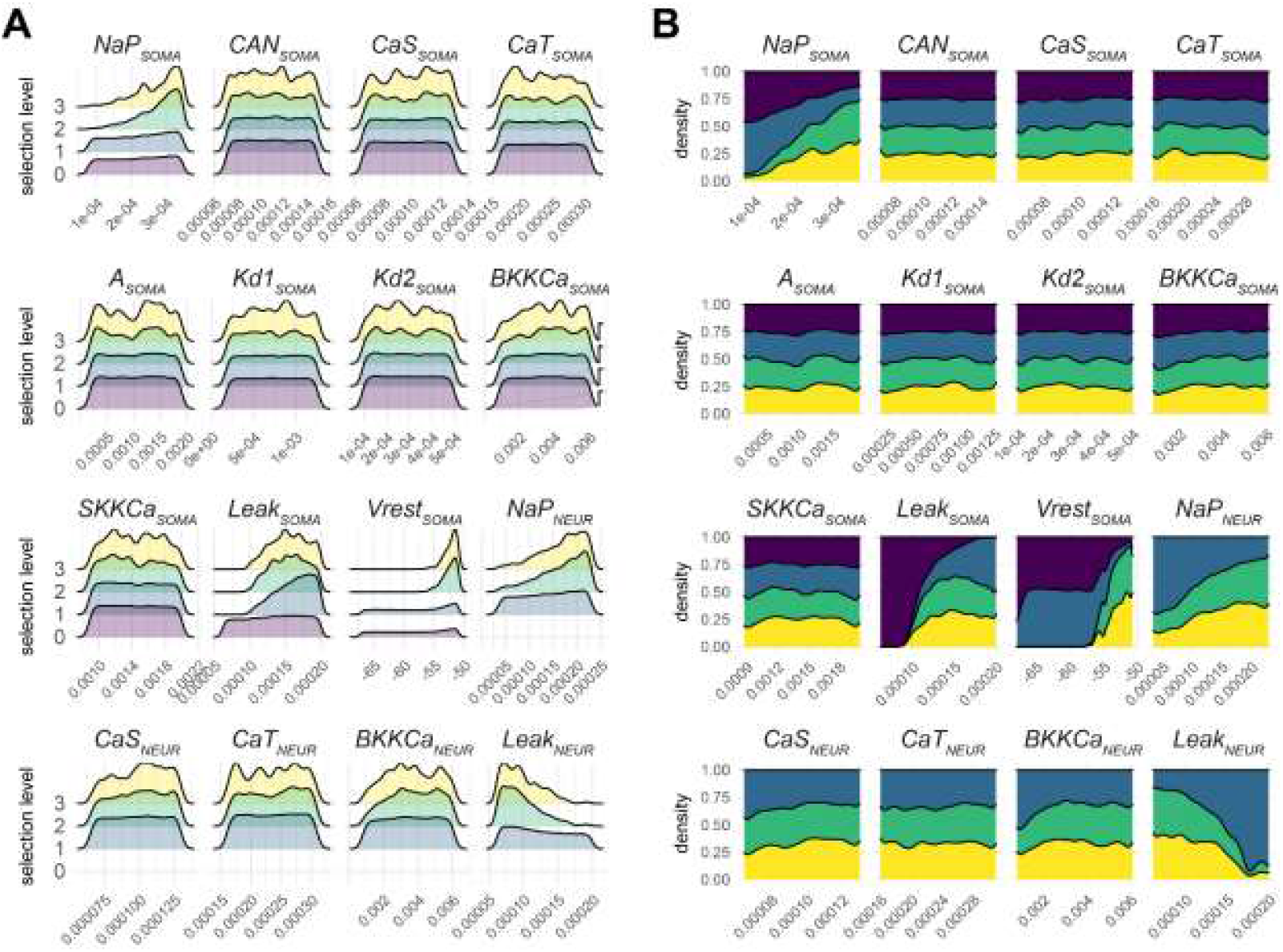
Conductance distributions after selection of soma and neurite model conductances. A.) Distribution of the conductances of models spared each level of selection; those selected on by the next level’s criteria. Each selection level (0, 1, 2, 3) represents the conductances that passed that level with the level 0 being those that were initially generated and level 3 being those which satisfied all criteria. Each level’s density plot is scaled independently. Because selection level 0 is performed on isolated somata, there are no conductances represented at this level for the neurite compartment. B.) Stacked density plots showing the subset of filtered at each level of selection. Four different levels of selection are shown (purple = 0, blue = 1, green = 2, yellow = 3). Each selection level (0, 1, 2, 3) represents the conductances of models that passed that level of selection, but not the subsequent – with the exception of level 3, which shows those that were preserved through all levels of selection. This indicates for a given conductance value which level of selection is most frequent.

#### Conductance parameter differences larger between strongly gap-junction coupled cells

Since the gap junction coupling between 4 and 5 is considerably stronger than between either of those and 3, we hypothesized that the network would be able to support a larger variation in conductances between 4 and 5 compared to the same between either of those and LC 3. To test for this, we estimated the Euclidean distance between the parameter sets of each of the three cells. Since LCs 4 and 5 are tightly coupled, we averaged the conductances between LCs 4 and 5, and then estimated the Euclidean distance between that average parameter set and that of LC3, for each network. These were then averaged across all the networks. This yielded the following when averaged across all the networks: the Euclidean distance between parameter sets of LC 4 and LC5 was 1.7689 (units?) and between averaged LCs4&5 and LC3 was 0.0148. The same calculations for LCs 1, 2 and 3 revealed the following numbers: the Euclidean distance between parameter sets of LC 1 and LC 2 was 2.0329 and between average of LCs1&2 and LC3 was 0.0155.

### Intrinsic conductance covariations in the network model

Biological studies have suggested that it is important and/or necessary for pairs of “modules” of conductance to work in concert to control appropriate physiological output. Therefore, we looked for relationships among all the membrane conductances in our model – across two compartments (soma and neurite) – at all three levels of selection.

Figure 5D describes membrane conductance relationships in the soma. Correlograms (Figure 5D, top) demonstrate that there is a general strengthening of conductance correlations [as evidenced by increasing absolute value of rho(ρ)-value] across the levels of selection. To better visualize individual relationships, we have plotted each pairwise correlation coefficient at each level of selection as a bar plot matrix (Figure 5D, bottom). We see several correlations that become apparent as network level selection of model neurons progresses. However, overall, the rho-values for any correlation remain relatively low, with the highest correlation coefficient approaching 0.2. Nevertheless, 5 conductance relationships emerge across the selection levels. The most substantial relationship is seen for Leak versus NaP in the soma (Figure 5D), and this emerges as a notable positive correlation. Two other positive correlations of some note are detected in the soma (Figure 5D): SKKCa versus Kd2, and SKKCa versus Leak. Furthermore, three negative correlations become apparent in the soma as well: CaT versus A, CaT versus BKKCa, and Leak versus A, and Bkkca and leak.

**Figure 5.**
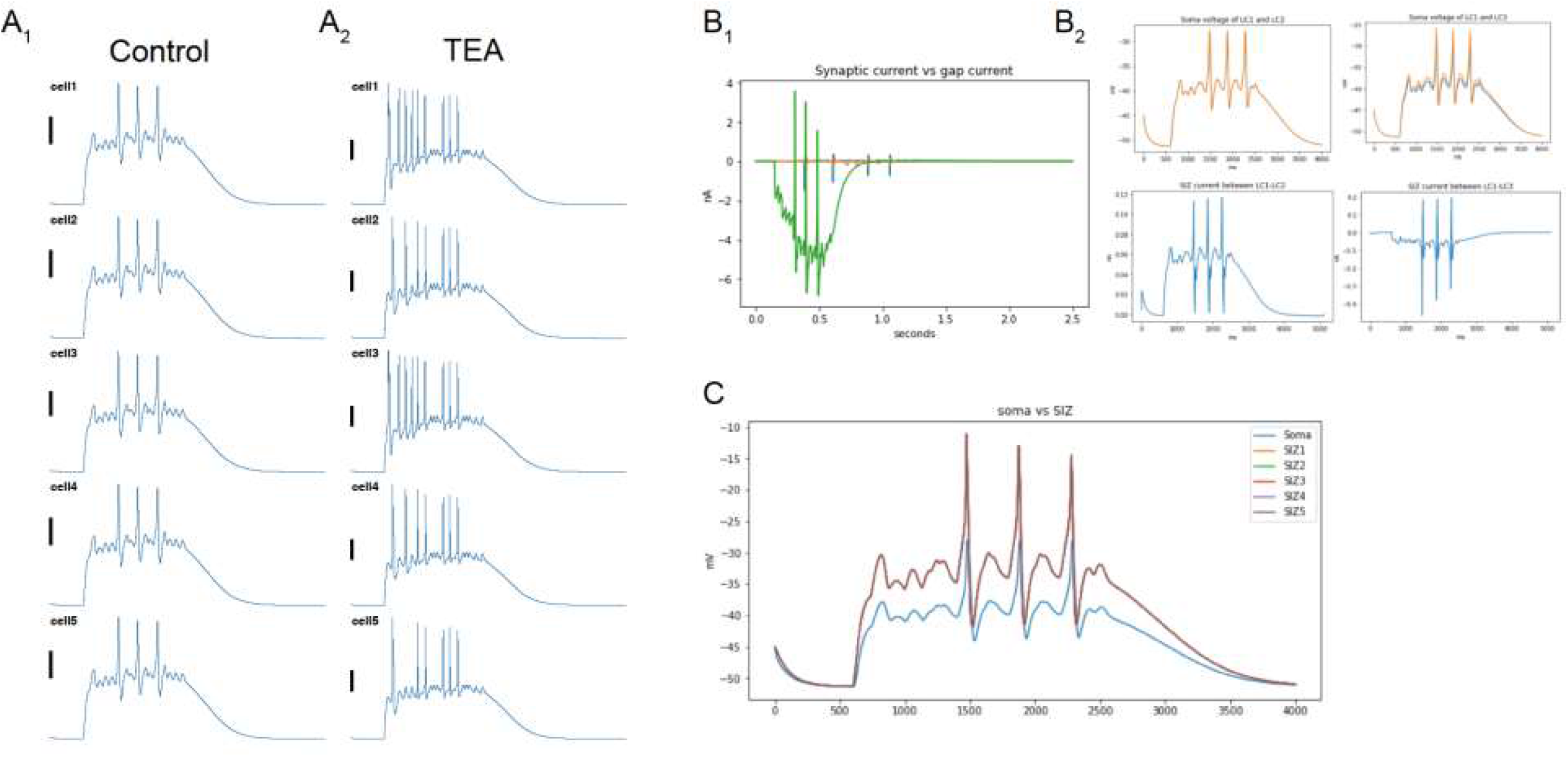
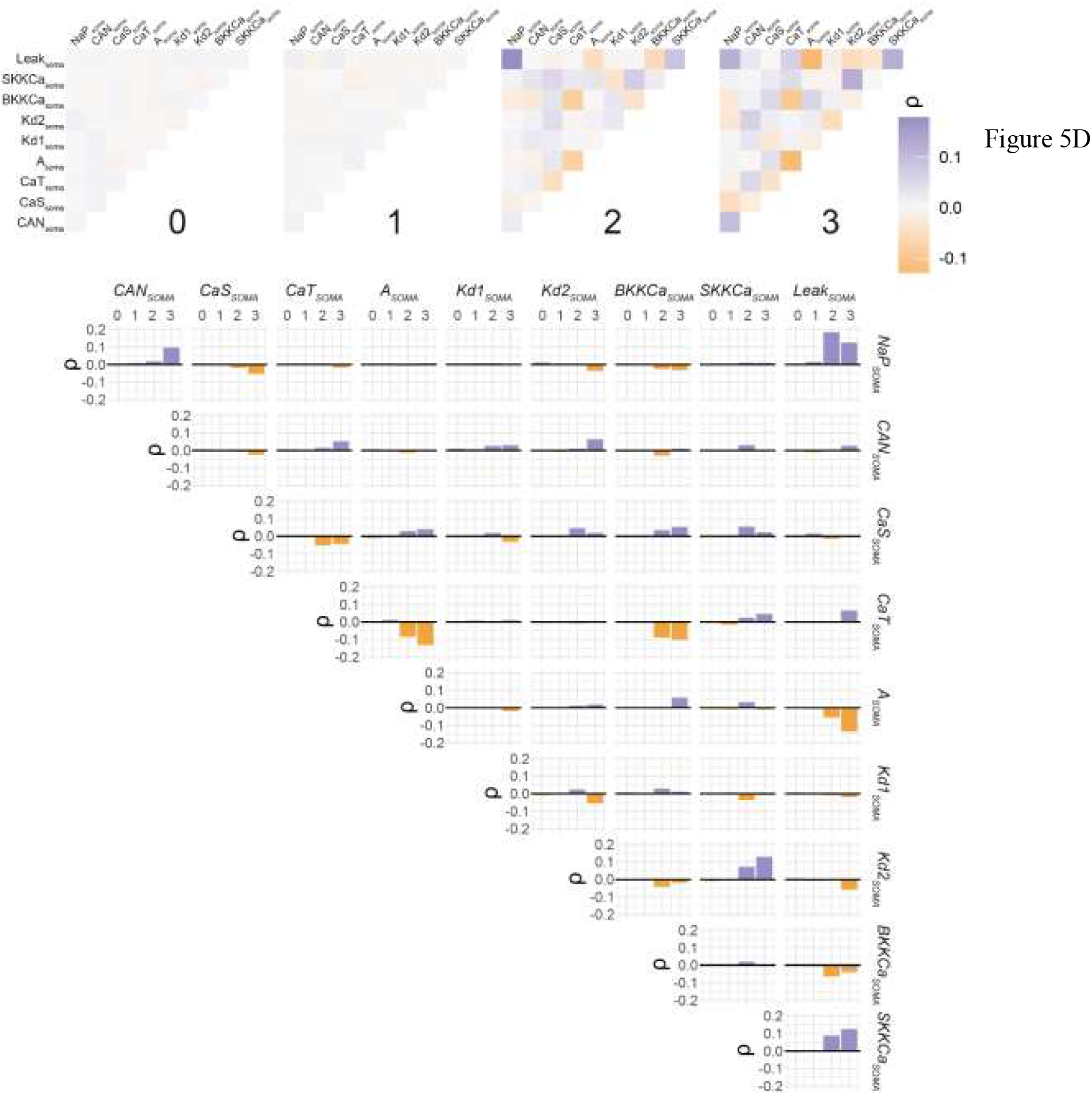
Conductance correlations in the soma compartment of model neurons across selection levels. TOP) Correlograms for four levels of selection (0-3) conductances in the somatic compartment of selected model neurons. Each pairwise correlation was calculated using Spearman’s correlation and reported as rho values. These plots demonstrate that correlations become more pronounced across subsequent levels of selection. BOTTOM) Bar plot showing the rho-value of each pairwise correlation across levels of selection. These are the same data plotted in the top row but allow for more precise determination of the more pronounced correlations.

Figure 6 describes the same conductance relationship analyses for the neurite compartment. Again, we can see relationships emerging and strengthening across the levels of selection (Figure 6, top). Two relationships appear more prominently in the neurite: a positive correlation between Leak and NaP, and a negative correlation between the two calcium conductances – CaS and CaT (Figure 6, bottom). Of note, Leak and NaP emerges as the strongest relationship in both the neurite and the soma compartments.

**Figure 6.**
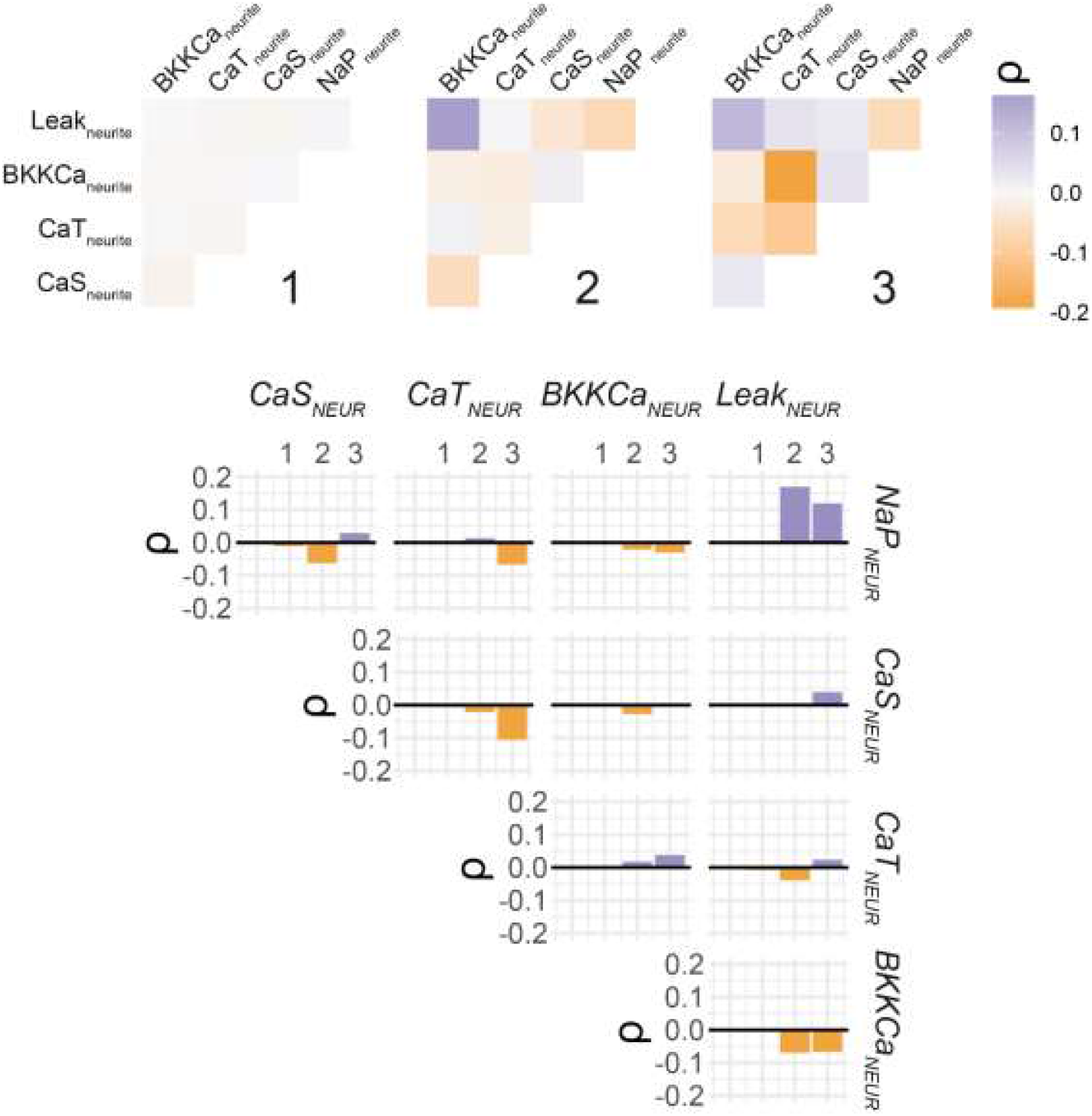
Conductance correlations in the neurite compartment of model neurons across selection levels. TOP) Correlograms for three levels of selection (1-3) conductances in the neurite compartment of selected model neurons as described in the previous figure. BOTTOM) Bar plot showing the rho-value of each pairwise correlation across levels of selection. These are the same data plotted in the top row but allow for more precise determination of the more pronounced correlations.

Finally, we comprehensively compare conductance relationships *across* compartments, to determine whether there may be co-regulation of membrane conductances between the soma and neurite. Figure 3 shows a comprehensive view of the conductance data in both compartments at selection level 3. In this visualization, we can see the distribution of each conductance along the diagonal, as well as the raw scatterplots of the 295 neurons at this level of selection. In addition to visualizing the raw data for the relationships in Figures 5D and 6, we see two strong relationships emerge across compartments: NaP_SOMA_ versus NaP_NEURITE_, and Leak_SOMA_ versus Leak_NEURITE_.

### Ion Channel mRNA correlations relative to model relationships

To determine whether the relationships seen among model neuron conductances may have independently arisen as necessary for appropriate output, we wanted to compare these results with a biological data set. Because it is difficult or impossible to comprehensively measure membrane conductances in biological neurons, we performed an analysis on levels of the mRNAs that encode the channels most directly responsible for membrane conductances that are represented in our model neurons. Using single-cell qPCR, we quantified 12 different channel mRNAs from 40 individual crab LC motor neurons (Figure 7).

**Figure 7.**
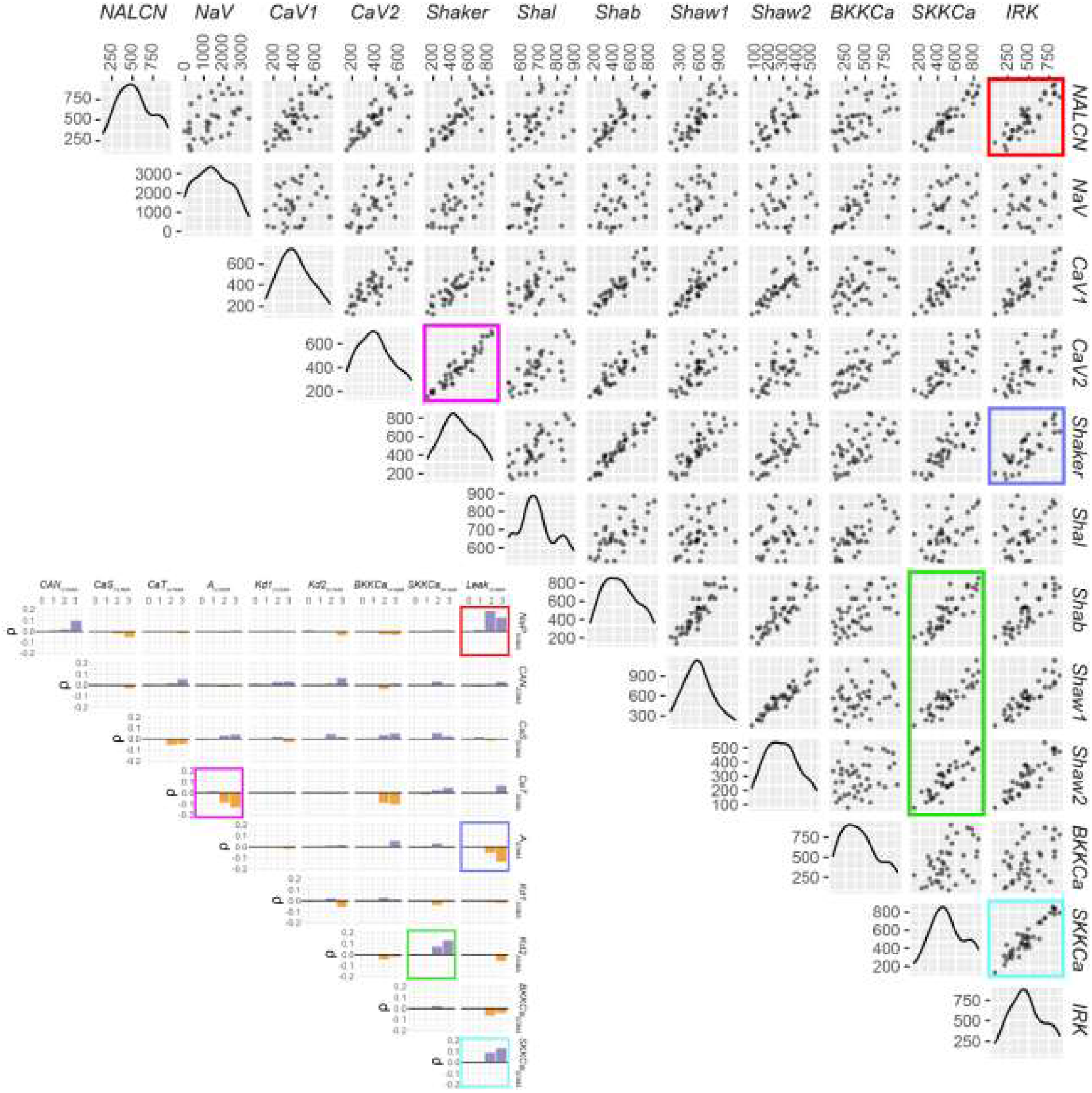
Scatterplots for pairwise relationships among channel mRNAs in biological LC motor neurons. Each dot represents a single model and its values for a given pair of channel mRNAs. Along the diagonal are curves that represent the distribution of values for a given conductance as keyed along the top axis. Stronger correlations that were identified in selection levels 2 and 3 in the soma of model neurons are noted by the colored box corresponding to the channel mRNAs most likely to encode those conductances. The inset shows the correlation bar plot from Figure X with the conductance relationships corresponding to a given pair of channel mRNAs color coded to match the mRNA plot.

While we are unable to disentangle mRNAs for channels that will be localized to the soma versus those that will be in the neurite, we feel the mRNA data are best compared with the somatic conductances in the model. The channel mRNAs reveal some very intriguing relationships. First, there is widespread correlation among mRNAs, and of considerably high correlation coefficients (Figure 7). Further, every correlation is a positive correlation. However, some of the most profound relationships bear striking similarity to those seen in the model. For example, *NALCN* and *IRK* are putatively related to the model conductances NaP and Leak respectively. NaP and Leak was the strongest correlation seen in the soma compartment and is a very tight correlation in biological cell mRNA as well (Figure 7). Furthermore, the four other strongest correlations found in the soma compartment of the model (CaT/A, Leak/A, SKKCa/Kd2, and SKKCa/Leak) reflect some of the strongest mRNA correlations in their biological channel counter parts as well (see Figure 8; *CaV2/Shaker, IRK/Shaker, SKKCa/Shab-Shaw1-Shaw2*, and *SKKCa/IRK* respectively). Thus, while the overall quantitative level of correlation differs, it is striking that the most tightly correlated mRNA relationships correspond well to the strongest model conductance correlations in the soma.

## DISCUSSION

We developed morphologically enhanced biophysical model of the LC of *Cancer Borealis* that included a new, hitherto unreported compartment SIZ that receives the synaptic input, and is informed completely by measured properties of the same cell. The model was used to investigate the potential role of membrane conductances in the neurite compartment of these cells. Furthermore, we engaged in a selection regime based on biologically relevant network activity that simultaneously selects five motor neurons for inclusion in our final cell population. Using network level output as selection criteria generated a population of 295 LC motor neurons in which we could further characterize relationships among membrane conductances in multiple compartments. In doing so, we identified correlations among ionic currents that are strongly reminiscent of channel mRNA relationships seen in a biological population of neurons. This is consistent with the hypothesis that neurons actively co-regulate membrane conductances to generate and maintain appropriate network output and suggests that these constraints may exist at a higher order network level rather than at the level of individual neuron properties.

### An intact model network generated from experimental data

Previous iterations of computational models of the crustacean cardiac ganglion [15; 16] relied on experimental data from multiple organs (e.g., stomatogastric ganglion) and species (e.g., lobster) for passive properties and conductance ranges. For the present model, the experimental data were obtained directly from the LCs of intact networks recorded from the Schulz lab, as well as SIZ recordings to first match our model with the known data, and then to explore the functional characteristics in search of an explanation for the preservation of output across a wide variation range. We started with a prediction of the intact single cell morphology that integrates information about structure and proposed a methodology to validate it. Specifically, although an SIZ has been conjectured as the source of SC input into the cardiac ganglion, reported single cell models have not explicitly considered such morphology. For instance, models of single LCs have typically considered only the soma compartment and possibly a neurite attached to it. To validate our enhanced three-compartmental model that includes an SIZ, we estimated the SC frequency ranges from intact recordings and then found that a model that accounted for the biological data was able to successfully reproduce intact single cell responses (Figs.2G)

### Biologically realistic framework for predictions single cell- and network-levels

Our new LC single cell model structure provided a framework to integrate experimental data related to LC input and output and explore underlying mechanisms. Specifically, the input to an LC arrived via synapses at its SIZ in the form of spike trains from the SCs. The range of frequency and temporal profiles of the SC spike trains followed experimental data. The output was the membrane potential fluctuation of the LC soma that in turn controlled the synchronization of spikes at the gap-junction coupled SIZ compartments (Fig. 1A). Using a three-stage rejection protocol to select the conductances in an unbiased manner, the procedure predicted that active conductances were necessary in the neurite to reproduce experimental input-output data for an LC.

To illustrate how a single-cell model is used to discover underlying mechanisms, recall that the LC model with soma with a short passive dendrite (to model the ligated neurite) was developed in stage 1 using experimental data from ligated LCs [22]. Using this model with a longer passive dendrite in stage 2 revealed that excessive leak in the passive neurite resulted in a reduction in depolarization of the soma membrane potential with spikes in the SIZ. This logic led to the consideration of Nap and/or CaS and CaT channels to counter the leakage. As shown in results, I_Nap and not I_CaS or I_CaT helped offset the effects of leak current in the neurite (Fig. 2B,D). The framework can be used to probe deeper underlying mechanisms. For instance, analysis of the time constants of the currents to probe how they accomplish this function, revealed that I_Nap had a time constant that was at least four-fold lower than that of I_CaT and at least 14-fold lower than that of I_CaS in the -50 to -20 mV range (Fig. 7). Since the membrane voltage of the soma never exceeded 0 mV, the time constant (during all model runs) of the I_CaT was always at least twice as large as that of I_Nap, and that of I_CaS was always larger than of I_CaT. However, I_CaT and I_CaS were continuously active and strengthened I_BKKCa as seen by comparing the traces in Figs. 2D, E.

Although the framework is designed to study both single cell and network level dynamics, the focus presently was largely on single-cell studies with only the issue of synchrony across LCs being considered at the network level. However, the framework can be readily used for network level studies. Some that we envisage in the future include potential co-variations of the synaptic or gap-junction conductances on the SIZs with the intrinsic conductances reported here.

### Emergent conductance correlations in the multicompartmental network model

One of the key hypotheses that this work tests is whether network level selection criteria will result in a population of models in which conductance correlations emerge. The collective literature in two similar crustacean networks (stomatogastric and cardiac ganglia) are somewhat inconsistent in this result. Previous computational models in the cardiac ganglion LCs from our group [15; 16]with a similar rejection sampling approach yielded strong correlations in the LC soma among two pairs of conductances: CaS-A and CaT-Kd. These data reveal that such relationships can emerge naturally from a selection process focused on output characteristics informed by biological data. However, those results were limited to models of the soma only and based on a different species and different mode of activity (driver potentials) than the cells modeled in our study. Further, there was relatively little biological data available at the time for those experiments, and so ultimately our interpretation of these first models is that such relationships are theoretically possible to detect and quantify – but further study was needed, including more thorough grounding in biological data as well as model neurons that better reflect the morphological complexity (i.e., multiple compartments) of biological neurons.

Conversely, a thorough and extensive analysis of a multicompartment model employing a large population of LP neurons from the crustacean stomatogastric ganglion yielded a different outcome. Taylor et al. [14] utilized a large population selection approach, in a multicompartment model, and selected based on output characteristics that focused both on the single cell excitability as well as some features reminiscent of network level function (i.e., output as a result of synaptic input currents). In this study, they found only weak correlations among conductances – both within and across the compartments [14]. Thus, our current work employs a much more similar approach to Taylor et al., but in the system originally modeled by previous members of our group – the cardiac motor neurons – in which we had seen such correlations. When we combined a multicompartmental model, far more extensive first-hand biological data, and a more developed network level selection process, our results in this study were largely consistent with those of Taylor et al. [14]. That is, while we could detect some emergent correlations within and across model neuron compartments, these were overall weaker correlations (rho-values between -0.2 and +0.2).

There are (at least) two overall interpretations of these results. First, that the intrinsic conductance correlations found in biological neurons across a wide range of nervous systems are not fundamentally *necessary* to generate baseline functional output of neurons. In other words, while these relationships may confer some adaptive advantage to neuron and network stability, and implicate compensatory relationships involved in homeostatic regulation, they are not in and of themselves fundamental to the solutions capable of producing a given output of a neuron in a network. However, we also interpret these results from a cautionary perspective. Taken together, the work of the Nair Lab shows that conductance correlations emerge in more narrowly constrained models with clear input-output relationships [15; 16]. However, as we add more free parameters to the system for which we have less knowledge of biological constraints – in this case multiple compartments and conductances therein, as well as synaptic inputs and network connectivity – we may lose the ability to detect fundamental relationships in biological neurons. If this is the case, then we predict that as more complex models become better informed by biological data, the possibility to recapitulate and interrogate these relationships may be more robust.

A more generous interpretation of the correlations found in the model would be that it is significant that such relationships can be detected at all given the conditions of the model experiment. Given that we had no biological data from which to inform the neurite compartment modeling, that the input-output relationships of LCs across individuals can be highly variable in biology, and that the network connectivity and synaptic drive from the pacemaker neurons had to be entirely inferred from the literature, we might predict that such levels of uncertainty would make it nearly impossible to expect biological relationships to emerge. Yet even though the correlation coefficients are somewhat weak, there is a clear constraint on conductances that emerge towards their correlated levels and several relationships are detectable as “signal above the noise.” Most provocative is the fact that the model conductance relationships we detect clearly are reflected as some of the most strongly correlated biological relationships at the level of channel mRNAs. While it would be inappropriate to overinterpret such disparate modalities of data (biological channel mRNAs are a long way removed from model membrane conductances), this provides some encouragement that better biological constraints to inform models going forward may recapitulate more strongly the relationships seen in biology. This will further allow computational modeling to be a critical test bed for exploring the nature of these relationships and how they influence output and stability in neural networks.

## CONCLUSION

We utilized a novel three-compartmental biophysical model of an LC that is morphologically realistic and includes provision for inputs from the SCs. The proposed single cell model facilitated incorporation of additional experimental observations related to both the SIZ compartment responses and to the presence of active conductances in the neurite compartment. Furthermore, the overall network model provided a framework to integrate this single cell information into a network and explore how it impacted and reproduced experimental observations at the network level. The model provided novel predictions of the differential roles of conductances in the neurite and the soma, and insights into the role of specific current channels in the neurite. The model also reproduced the varied responses seen experimentally and predicted the calcium currents in the neurite to be the underlying cause. Finally, we investigated whether conductance relationships would emerge from the selection process that would provide insight into the biological function of these interactions, as well as allow us to make inferences about the fundamental nature of such relationships in biological neurons. While we did detect some correlations among conductances within and across compartments, these were overall weaker relative to our previous work and that reported in the biological literature. We suggest that either such conductance relationships are not fundamentally necessary to generate a given output, or that much greater constraints on the free model parameters are needed to recapitulate these biological relationships. Overall, these predictions and the reasons why the LCs exhibit varied TEA responses are topics for future research.

## List of Figures

Figure 1. **Biological realistic model of intact LC reproduces experimental dat**a. - still being updated

Figure 2. **Model predicts presence and roles of active conductances in neurite**. – finalized

Figure 3. **Scatterplots for pairwise relationships among soma, neurite, and across compartment conductances in models after selection level 3**.

Figure 4. **Conductance distributions after selection of soma and neurite model conductances**. A.) Distribution of the conductances of models spared each level of selection; those selected on by the next level’s criteria. Each selection level (0, 1, 2, 3) represents the conductances that passed that level with the level 0 being those that were initially generated and level 3 being those which satisfied all criteria. Each level’s density plot is scaled independently. Because selection level 0 is performed on isolated somata, there are no conductances represented at this level for the neurite compartment. B.) Stacked density plots showing the subset of filtered at each level of selection. Four different levels of selection are shown (purple = 0, blue = 1, green = 2, yellow = 3). Each selection level (0, 1, 2, 3) represents the conductances of models that passed that level of selection, but not the subsequent – except for level 3, which shows those that were preserved through all levels of selection. This indicates for a given conductance value which level of selection is most frequent.

Figure 5. **Conductance correlations in the soma compartment of model neurons across selection levels**. TOP) Correlograms for four levels of selection (0-3) conductances in the somatic compartment of selected model neurons. Each pairwise correlation was calculated using Spearman’s correlation and reported as rho values. These plots demonstrate that correlations become more pronounced across subsequent levels of selection. BOTTOM) Bar plot showing the rho-value of each pairwise correlation across levels of selection. These are the same data plotted in the top row but allow for more precise determination of the more pronounced correlations.

Figure 6. **Conductance correlations in the neurite compartment of model neurons across selection levels**. TOP) Correlograms for three levels of selection (1-3) conductances in the neurite compartment of selected model neurons as described in the previous figure. BOTTOM) Bar plot showing the rho-value of each pairwise correlation across levels of selection. These are the same data plotted in the top row but allow for more precise determination of the more pronounced correlations.

Figure 7. **Scatterplots for pairwise relationships among channel mRNAs in biological LC motor neurons**. Each dot represents a single model and its values for a given pair of channel mRNAs. Along the diagonal are curves that represent the distribution of values for a given conductance as keyed along the top axis. Stronger correlations that were identified in selection levels 2 and 3 in the soma of model neurons are noted by the colored box corresponding to the channel mRNAs most likely to encode those conductances. The inset shows the correlation bar plot from Figure X with the conductance relationships corresponding to a given pair of channel mRNAs color coded to match the mRNA plot.

## METHODS

### Experimental data to constrain single cell and network models

The biological data used to constrain both the LC model parameters (e.g., membrane currents) and outputs (of both ligated and intact LCs) were collected under the auspices of previously published work as follows.

Membrane currents were made in two-electrode voltage clamp while the network activity was silenced either with tetrodotoxin (TTX) or by severing the CG nerve trunk to remove the small cell (SC) inputs. The inward currents I_CaS_, I_CaT_, I_NaP_, and I_CAN_ were based on recordings and data as described in Ransdell et al. [1]. The outward currents I_A_, I_Kd_, I_BKKCa_ were based on recordings made in Ransdell et al. [2]. No biological characterization of SKKCa has been performed in crabs, and this work carries over SKKCa model currents as described in our previous CG modeling efforts [3]. Intracellular voltage follower recordings of ongoing network activity were made in all of the above studies, and from these we generated the biological parameters to constrain model network output.

Synaptic inputs (chemical) and connections (electrical) were characterized from these recordings as well. EPSPs were characterized by measuring the amplitude and time constant characteristics from intracellular LC recordings in intact networks. Single SC action potentials that yielded clear (non-summating) EPSPs were used to generate a population of post-synaptic potential measurements that constrained the chemical synapse inputs. Electrical coupling was measured directly in two-electrode current clamp as described in Lane et al. [3]

To characterize isolated LC soma responses with and without TEA, we used a current clamp protocol designed to emulate SC synaptic inputs. These chemical synapse stimulus protocols are described in detail in Ransdell et al. [1].

Finally, mRNA levels for ion channels were taken from previously published work by Northcutt et al. [4] and analyzed and formatted for use towards the experimental goals in this study.

### Development of biophysical single cell models

The single cell model had three compartments: soma, neurite (neu) and spike-initiation-zone (SIZ). The soma compartment had a length of 120 µm and a diameter of 90 µm, and contained 9 currents. The neurite had a length of 1380 µm, and a diameter of 12 µm, and contained 5 currents. The SIZ had a length of 108 µm, and a diameter of 20 µm, and contained 3 currents. The Na and K channels in the SIZ were given fixed conductances of 0.2 and 0.4 S/cm^2^, respectively, and we assumed a specific capacitance of 1.5 μF/cm^2^ for all three structures. The model for currents for each compartment followed the Hodgkin-Huxley equation formulation (Eqn. 1)

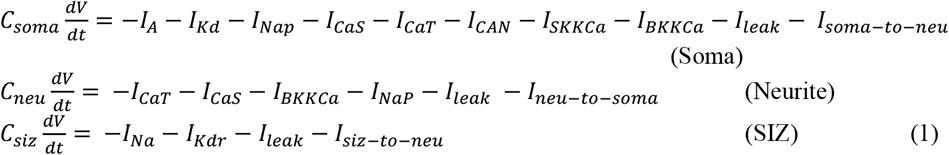

where the currents on the right-hand side of the first equation are: A-type potassium (I_A_), delayed rectifier (I_Kd_), persistent sodium (I_Nap_), slow persistent calcium (I_CaS_), transient calcium (I_CaT_), calcium-dependent non-selective cation (I_CAN_), twSo calcium-dependent potassium currents (I_SKKCa_ and I_BKKCa_), leak (I_Leak_) and the injected current (I_inj_). The individual currents were modeled as *I*_*c*_ = *g*_*max,c*_*m*^*p*^*h*^*q*^(*V* − *E*_*c*_), where *g*_*max,c*_ is its maximal conductance, *m* its activation variable (with exponent *p*), *h* its inactivation variable (with exponent *q*), and *E*_*c*_ its reversal potential (a similar equation is used for the synaptic current but without *m* and *h*). The kinetic equation for each of the gating functions *x* (*m* or *h*) takes the form

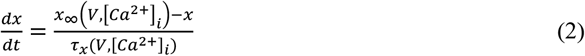

where *x*_∞_ is the steady state gating voltage- and/or Ca^2+^- dependent gating variable and *τ*_*x*_ is the voltage**-** and/or Ca^2+^**-** dependent time constant. The equations for the active channels in the soma compartment were fit using biological recordings for these currents from our Lab from the cardiac ganglion of *Cancer borealis*. These currents were fit as follows: Voltage clamp data obtained with Clampfit were imported into MATLAB and fit using the MATLAB curve-fitting toolbox. Current data were converted to conductance data by dividing by (V_m_ – E_Rev_), where E_Rev_ was as follows: E_Na =_ +55 mV, E_K_ = -80 mV, E_Ca_ = +45 mV, E_Leak_ was chosen uniformly randomly from [-67.1,50.6], and E_CAN_ = -30 mV. The time axis was adjusted to start from 0 for the beginning of the clamp. The following parametrization was used:

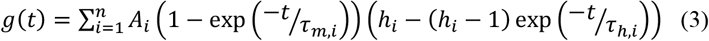

In this equation, A_i_ = G_i,max_ × m_i_ was the maximal conductance of the current i multiplied by its voltage-dependent steady-state activation (m_i_), h_i_ was the steady-state inactivation value, and τ_m,i_ and τ_h,i_ were the time constants with which activation and inactivation reached steady-state, respectively. This fitting procedure assumed that ionic currents were completely deactivated m=0 and deinactivated (h=1) prior to the onset of the voltage clamp. This was fit to each trace in voltage clamp experiment, giving values of each of the four parameters for each test clamp voltage (V_c_). These values were then fit for each current as functions of V_c_ using the general forms as stated below. This procedure yielded equations for the currents recorded in voltage clamp that could be used in simulations according to the Hodgkin-Huxley formalism.

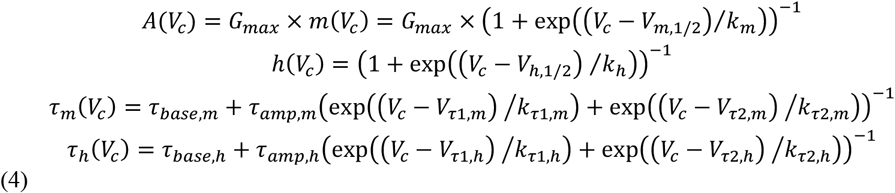

All the maximal conductances (G_i,max_) were in µS, time constants in ms and voltages in mV (4)

#### Calcium dynamics

Intracellular calcium modulates the conductances of the calcium-activated potassium currents (BKKCa and SKKCa), calcium-activated nonselective cation current (CAN) and also influences the magnitude of the inward calcium current in the LC [5]. A calcium pool was modeled in the LC with its concentration governed by the first-order dynamics [6; 7] below:

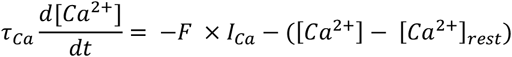

where F = 0.256 μM/nA is the constant specifying the amount of calcium influx that results per unit (nanoampere) inward calcium current; τ_Ca_ represents the calcium removal rate from the pool; and [Ca^2+^]_rest_ = 0.5 μM. Voltage-clamp experiments of the calcium current in the our lab showed the calcium buffering time constant to be around 690 ms (τ_Ca_).

### Searching for viable biophysical LC neurons within the model parameter space

We used a three-stage rejection protocol to select viable networks with five distinct LC cells, with each LC satisfying biological single cell and current injection responses. For the single cell model search, we started with a set 100,000 three-compartment cell models with the 14 conductances and the small cell frequency selected randomly from uniform distributions of values between their respective minimum and maximums given in table 3. The SC frequency range of 16-32 Hz was determined from SC recordings from our lab (see Supplementary Materials, table 1). The recordings showed that the combined small and medium spike frequency averages had minimum and maximum frequency of 16 Hz and 32 Hz, respectively.

In stage 1 of the rejection protocol, ligated soma (soma + neurite) models were tested for passive properties of resting membrane potential, time constant and input resistance. Cells for which these values were within ranges in table 4 were retained. In stage 2, SIZ was added to the passing cells and a synaptic input (spike train mimicking SC input) was provided. Based on biological recordings from our Lab, on average, the average values for an SC burst were as follows: period was 1000 ms, and the synaptic input frequency increase for 600 ms starting at 300 ms. The initial SC burst was for 300 ms, and this was followed with a higher frequency for 600 ms, i.e., till 900 ms, after which it fell back to the original frequency and terminated at 1000 ms. The synapse had a fixed gain of 0.0309 μS in LV2. The reversal potential was chosen as -15 mv, from STG studies. This was chosen from a range of values as the one that produced the most passing cells in the tested sample. Values were picked from a range between 0 -0.2 uS, since 0.2 uS restored synchrony in TEA conditions [3]. The frequency of SC input is randomly picked such that the maximum frequency of the input is chosen. For example, for the first iteration, a number between 16 and 17 is chosen, say 16.4, and the input at 300 -600 ms would be 40% of 16.4, then the for 600 ms would be 16.4 hz, then for 100ms would be 40% of 16.4 Hz. The current was delivered to the SIZ using Neuron’s VecStim. Events were delivered to VecStim from a csv which was generated based on the random SCfrequency chosen. Each cell was tested at each frequency interval between 16 and 32 Hz.i.e., a random frequency between 16-17, then a random frequency between 17-18, etc. Each network was also tested at each frequency interval. The reason for using randomly chosen intervals rather than fixed numbers was so that more frequencies would be used and the distribution of passing cells across frequencies would be more even. The network model uses gap junctions between LC1 and LC2 and LC4 and LC5, with gap junctions between all SIZs. The coupling conductance between the large cells is 0.65 μS, and is 0.067 μS between the SIZs. There is no recorded coupling conductance for SIZ, so this conductance was chosen based on what produced a passing network, however a previous study reported coupling conductances between the LC’s to be between 0.4-0.8 uS [3]. When perfused with TEA, the cells shows adaptive conductance changes to be between 0.8-1.2.

Both control and post-TEA case are considered and intact cells whose resulting membrane potential waveform characteristics were within the ranges shown in table 6 were retained. In a third stage, we assemble networks and select viable ones as described next.

### Determination of viable network models using the selected single cell models

First, we randomly selected five distinct LCs for the network from the pool of intact LCs that pass Stage 2. Then we connected an SIZ to the distal end of the neurite of each LC, and attached a synapse to the SIZ. Also, we connected the soma compartments of LCs 4 and 5 and of LCs 1 and 2 with separate gap junction values that were determined experimentally in our lab (Table 3). The SC pacemaker drive was delivered as a spike train to the five excitatory synapses on each SIZ. It was observed biologically that frequency of SC firing increases within the slow wave oscillation cycle of LCs. The SIZ compartment and the excitatory SC-SIZ synapse configuration was identical for each LC, i.e., we did not vary model parameters for these compartments and synapses.

Experimental TEA block was simulated by reducing the conductances G_BKKCa_, G_Kd_ and G_A_ by 97% in the LCs and neurite [1]. Synchrony scores were computed by using the R^2^ between the large cells 3 and 5. To determine the range for synchrony scores, we examined five experimental recordings for LC3 and LC5 (Unpublished data). Taking the first two minutes of TEA exposure (acute), we measured the R^2^ between LC3 and LC5 recordings. We chose the maximum of TEA synchrony to be the lowest control synchrony score minus 1.5 times the control interquartile range. The lowest control synchrony score was 0.9425, and the control interquartile range was 0.0318, therefore the maximum of TEA synchrony was taken to be 0.8948. The cells were considered desynchronized if the synchrony score was below 0.9425. For the synchrony score between LC3 and LC5, we considered anything below 0.89 to be asynchronous, and anything above 0.9425 to be demonstrating synchrony.

After performing control and TEA runs using these networks, in the third stage, we rejected networks that had waveform characteristics outside the ranges shown in Table 7. We rejected networks that showed excessive TEA synchrony between LC3 and LC5, or insufficient spikes per burst since neither behavior was observed in biological traces. This left xx networks that reproduced the biological trends and these were used in subsequent analyses to explore potential conductance changes that could restore network synchrony.

## REFERENCES

1. Marder E (2011) Variability, compensation, and modulation in neurons and circuits, Proc Natl Acad Sci U S A 108 Suppl 3, 15542-15548. PMC3176600.

2. Prinz AA, Billimoria CP, Marder E (2003) Alternative to hand-tuning conductance-based models: Construction and analysis of databases of model neurons, J Neurophysiol 90, 3998–4015.

3. Goaillard JM, Taylor AL, Schulz DJ, Marder E (2009) Functional consequences of animal-to-animal variation in circuit parameters, Nat Neurosci 12, 1424-1430. PMC2826985.

4. MacLean JN, Zhang Y, Goeritz ML, Casey R, Oliva R, Guckenheimer J, Harris-Warrick RM (2005) Activity-independent coregulation of ia and ih in rhythmically active neurons, J Neurophysiol 94, 3601–3617.

5. Khorkova O, Golowasch J (2007) Neuromodulators, not activity, control coordinated expression of ionic currents, The Journal of neuroscience : the official journal of the Society for Neuroscience 27, 8709–8718.

6. Schulz DJ, Goaillard J-M, Marder E (2006) Variable channel expression in identified single and electrically coupled neurons in different animals, Nat Neurosci 9, 356–362.

7. Schulz DJ, Goaillard J-M, Marder EE (2007) Quantitative expression profiling of identified neurons reveals cell-specific constraints on highly variable levels of gene expression, Proceedings of the National Academy of Sciences 104, 13187–13191.

8. Swensen AM, Bean BP (2005) Robustness of burst firing in dissociated purkinje neurons with acute or long-term reductions in sodium conductance, J Neurosci 25, 3509-3520. PMC6725377.

9. Marder E, Goaillard J-M (2006) Variability, compensation and homeostasis in neuron and network function, Nat Rev Neurosci 7, 563–574.

10. Goldman MS, Golowasch J, Marder E, Abbott LF (2001) Global structure, robustness, and modulation of neuronal models, J Neurosci 21, 5229–5238.

11. Tobin AE, Van Hooser SD, Calabrese RL (2006) Creation and reduction of a morphologically detailed model of a leech heart interneuron, J Neurophysiol 96, 2107–2120.

12. Olypher AV, Calabrese RL (2007) Using constraints on neuronal activity to reveal compensatory changes in neuronal parameters, J Neurophysiol 98, 3749–3758.

13. Gunay C, Edgerton JR, Jaeger D (2008) Channel density distributions explain spiking variability in the globus pallidus: A combined physiology and computer simulation database approach, J Neurosci 28, 7476–7491.

14. Taylor AL, Goaillard J-M, Marder E (2009) How multiple conductances determine electrophysiological properties in a multicompartment model, The Journal of Neuroscience 29, 5573–5586.

15. Ball JM, Franklin CC, Tobin A-E, Schulz DJ, Nair SS (2010) Coregulation of ion channel conductances preserves output in a computational model of a crustacean cardiac motor neuron, J Neurosci 30, 8637-8649. PMC4473856.

16. Franklin CC, Ball JM, Schulz DJ, Nair SS (2010) Generation and preservation of the slow underlying membrane potential oscillation in model bursting neurons, J Neurophysiol 104, 1589–1602.

17. Olypher AV, Prinz AA (2010) Geometry and dynamics of activity-dependent homeostatic regulation in neurons, J Comput Neurosci 28, 361–374.

18. Marder E, Taylor AL (2011) Multiple models to capture the variability in biological neurons and networks, Nat Neurosci 14, 133–138.

19. Lane BJ, Samarth P, Ransdell JL, Nair SS, Schulz DJ (2016) Synergistic plasticity of intrinsic conductance and electrical coupling restores synchrony in an intact motor network, eLife 5, e16879. PMC5026470.

20. Tobin A-E, Cruz-Bermúdez ND, Marder E, Schulz DJ (2009) Correlations in ion channel mrna in rhythmically active neurons, PLoS One 4, e6742.

21. O’Leary T, Williams AH, Franci A, Marder E (2014) Cell types, network homeostasis, and pathological compensation from a biologically plausible ion channel expression model, Neuron 82, 809–821.

22. Ransdell JL, Nair SS, Schulz DJ (2012) Rapid homeostatic plasticity of intrinsic excitability in a central pattern generator network stabilizes functional neural network output, J Neurosci 32, 9649–9658.

23. Cruz-Bermúdez ND, Marder E (2007) Multiple modulators act on the cardiac ganglion of the crab cancer borealis, J Exp Biol 210, 2873–2884.

24. Ransdell JL, Nair SS, Schulz DJ (2013) Neurons within the same network independently achieve conserved output by differentially balancing variable conductance magnitudes, J Neurosci 33, 9950–9956.

## REFERENCES

1. Ransdell JL, Nair SS, Schulz DJ (2013) Neurons within the same network independently achieve conserved output by differentially balancing variable conductance magnitudes, J Neurosci 33, 9950–9956.

2. Ransdell JL, Nair SS, Schulz DJ (2012) Rapid homeostatic plasticity of intrinsic excitability in a central pattern generator network stabilizes functional neural network output, J Neurosci 32, 9649–9658.

3. Lane BJ, Samarth P, Ransdell JL, Nair SS, Schulz DJ (2016) Synergistic plasticity of intrinsic conductance and electrical coupling restores synchrony in an intact motor network, eLife 5, e16879. PMC5026470.

4. Northcutt AJ, Kick DR, Otopalik AG, Goetz BM, Harris RM, Santin JM, Hofmann HA, Marder E, Schulz DJ (2019) Molecular profiling of single neurons of known identity in two ganglia from the crab cancer borealis, Proc Natl Acad Sci U S A 116, 26980–26990.

5. Tazaki K, Cooke I (1990) Characterization of ca current underlying burst formation in lobster cardiac ganglion motorneurons, J Neurophysiol 63.

6. Prinz AA, Billimoria CP, Marder E (2003) Alternative to hand-tuning conductance-based models: Construction and analysis of databases of model neurons, J Neurophysiol 90, 3998–4015.

7. Soto-Treviño C, Rabbah P, Marder E, Nadim F (2005) Computational model of electrically coupled, intrinsically distinct pacemaker neurons, J Neurophysiol 94, 590-604. PMC1941697.

